# Stress-impaired reward pathway promotes distinct feeding behavior patterns

**DOI:** 10.1101/2021.12.08.471524

**Authors:** Yusuke Fujioka, Kaori Kawai, Kuniyuki Endo, Minaka Ishibashi, Nobuyuki Iwade, Dilina Tuerde, Kozo Kaibuchi, Takayuki Yamashita, Akihiro Yamanaka, Masahisa Katsuno, Hirohisa Watanabe, Shinsuke Ishigaki, Gen Sobue

## Abstract

Psychosocial stress can impact feeding behavior outcomes. Although many studies have examined alterations to food intake, little is known about how stress affects feeding behavior patterns. To determine the impact of psychological stress on feeding behavior patterns, mice were subjected to various psychosocial stressors (social isolation, intermittent high-fat-diet, or physical restraint) prior to timed observations in a feeding arena that incorporated multiple bait loci. In addition, in vivo microdialysis was used to assess the effects of stressors on the reward system by measuring dopamine levels in the nucleus accumbens (NAcc) shell. Impaired feeding behavior patterns characterized by significant deviations in bait selection (i.e. fixated feeding) and prolonged periods of eating (i.e. protracted feeding) were observed in stressed mice relative to non-stressed controls. In addition to clear behavioral effects, the stressors also negatively impacted dopamine levels at the nucleus accumbens shell. Normalization of dopamine reversed the fixated feeding behavior, whereas specifically inhibiting neuronal activity in the dopaminergic neurons of the ventral tegmental area that project to the nucleus accumbens shell caused similar impairments in feeding. Given that the deviations were not consistently accompanied by changes in the amount of bait consumed, body weight, or metabolic factors, the qualitative effects of psychosocial stressors on feeding behavior likely reflect perturbations to a critical pathway in the mesolimbic dopamine system. These findings provide compelling evidence that aberrations in feeding behavior patterns can be developed as sensitive biomarkers of psychosocial stress and possibly a prodromal state of neuropsychiatric diseases.

**Significance Statement:** Feeding behavior can be affected by neuropsychiatric disorders including psychosocial stressors, and the evaluation of eating behavior was mainly based on food intake. However, it is speculated that not only food intake but also feeding behavior patterns can be affected in such disorders. The biological processes underlying the feeding behavior patterns have not been clarified yet. We found that aberrant feeding behaviors in mice characterized by fixated feeding were provoked by psychosocial stressors. The qualitative effects of psychosocial stressors on feeding behavior reflect perturbations in the mesolimbic dopamine system. These findings provide compelling evidence that aberrations in feeding behavior patterns can be developed as sensitive biomarkers of psychosocial stress and possibly a prodromal state of neuropsychiatric diseases.

## Introduction

Eating behaviors in humans and other animals are tightly regulated by diverse regulatory brain circuits, many of which overlap with the brain’s reward system (Saper et al., 2002; Morton et al., 2014; Rossi and Stuber, 2018). Quantitative aspects of feeding behaviors are defined by the degree of food intake and are mainly controlled by hypothalamic circuits and peripheral metabolic hormones in response to energy demands. In contrast, qualitative aspects of feeding behaviors are determined by feeding patterns. The biological processes underlying the feeding behavior patterns, however, remain to be fully clarified.

Feeding behavior patterns can be affected by any number of psychosocial stressors that are potentially reinforced by COVID-19 confinement (Bemanian et al., 2020; Marchitelli et al., 2020; Cecchetto et al., 2021; Francois et al., 2021) in addition to neuropsychiatric disorders such as autism spectrum disorders (ASD) and front temporal lobar degeneration (FTLD) (Ikeda et al., 2002; Ahmed et al., 2016; Bandini et al., 2017). Since deviations in food preference can be caused by aberrant reward processing (Adam and Epel, 2007; Alonso-Alonso et al., 2015), stressors may disrupt feeding behavior patterns via the reward system. Thus, assessing feeding behavior patterns could be useful when evaluating eating disorders and associated stress states; however, eating disorders have been diagnosed by criteria that are based on food intake (Erzegovesi and Bellodi, 2016), and our understanding of the mechanisms underlying how various stress conditions disrupt normal feeding behavior patterns is limited. In particular, quantifiable methods that facilitate the study of feeding behavior patterns are needed.

In the present study, we developed a real-time monitoring system in tandem with a regular diet to assess the spatial and temporal aspects of feeding behavior patterns that allows for more accurate detection of the relative physiological states of animals. Using the system, we found that different stress conditions induced alterations in the spatiotemporal feeding patterns which occurred independent of metabolic alterations but were strongly associated with impairments in the mesolimbic dopamine system.

## Materials and Methods

### Mice and stress models

To investigate the effects of isolation stress on feeding behaviors, we randomly assigned wild-type C57BL/6J strain mice to either an experimental group that housed solitary mice or a control group. For the control group, mice were housed 4 mice together according to standard animal care protocols. For the experimental group, mice were housed alone for 7 days prior to the behavioral experiments and *in vivo* microdialysis. Social isolation is an environmental stressor that can alter feeding behaviors, but does not necessarily affect body weight (Yamada et al., 2015). It is thus a good model for elucidating the mechanisms underlying altered feeding behavior patterns in response to stress that is not accompanied by alterations in food intake and body weight.

Temporally limited access to palatable diets is a common method to induce binge eating behaviors in rodents (Avena and Bocarsly, 2012). As a stress model for increased food intake and body weight, we established an intermittent high-fat diet (HFD) model by randomly assigning 6-week-old, inbred male mice (C57BL/6J strain) to either an intermittent HFD group or a control group. The mice were placed in cages with free access for both standard chow diet and drinking water except for HFD treatment. The intermittent HFD group could access HFD (Test Diet 58Y1, PMI Nutrition International, KS, USA; 23.1% protein, 34.9% fat, 25.9% carbohydrate, and 6.5% fiber) for 2 h every other day, whereas ad libitum supply of standard chow (CE2, CLEA, Tokyo, Japan; 24.8% protein, 4.6% fat, 49.9% NEF, and 4.65% fiber) was used for the control group mice. After 2 weeks of HFD treatment, real-time monitoring of feeding behaviors and *in vivo* microdialysis of mice from both groups were performed.

To investigate the effects of physical stress on feeding behaviors accompanied by reduced food intake and body weight, inbred C57BL/6J male mice were randomly assigned to either a restrained group or an unimpeded movement control group. Six-week-old mice in the restrained group were completely immobilized using disposable mouse DecapiCone restrainers (MDC-200, Braintree Scientific, Inc. MA, US) for 2 h over 5 consecutive days prior to assessing feeding behaviors or *in vivo* microdialysis analyses (Fig. 1*A*). Detailed protocols have been previously described (Besnard et al., 2018).

**Figure 1.**
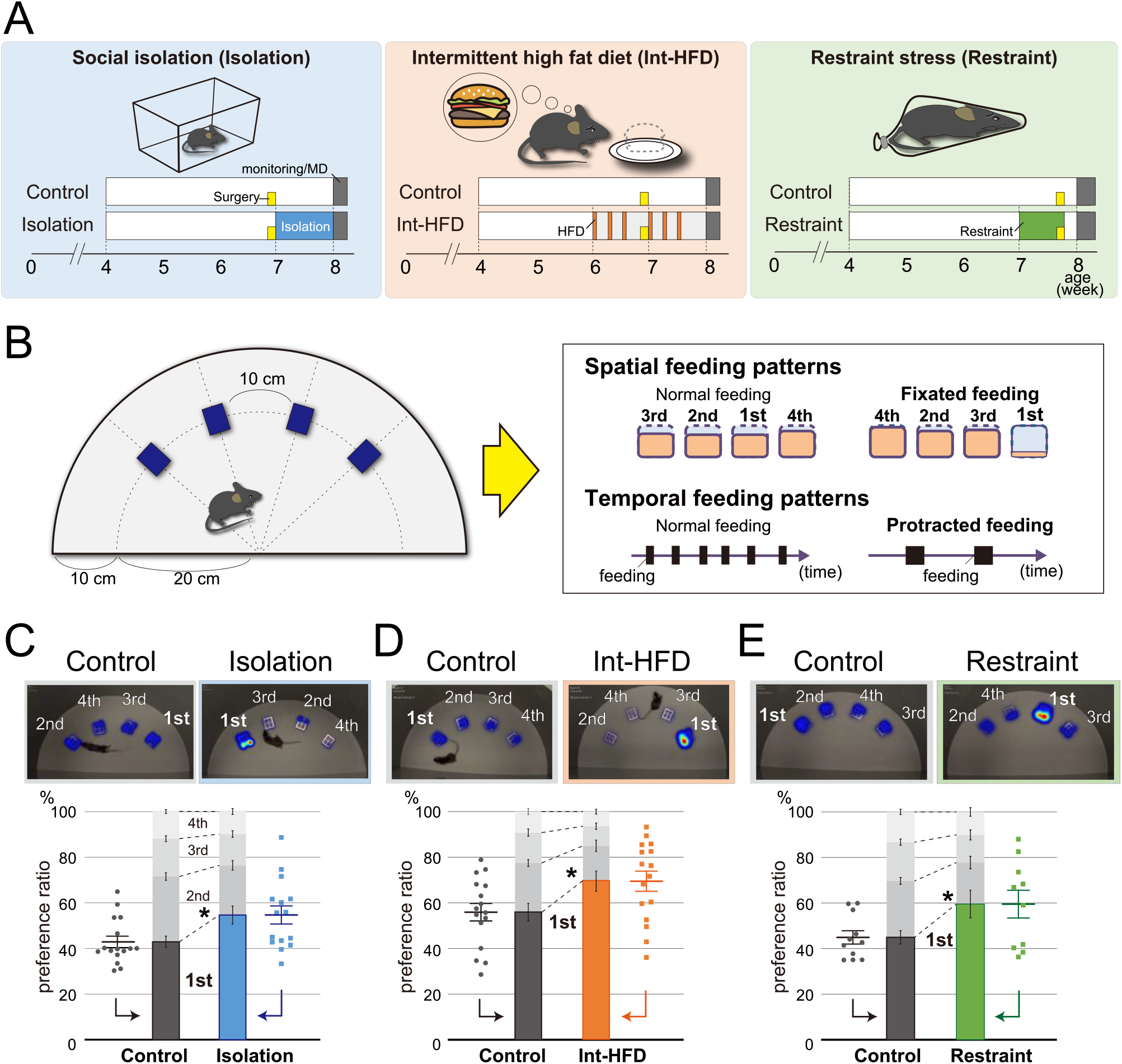
Psychosocial stresses such as social isolation, intermittent high-fat diet, and physical restraint cause aberrant feeding behavior patterns characterized by fixated feeding. ***A***, Study paradigms for social isolation (Isolation), intermittent high fat diet (Int-HFD), and restraint stress (Restraint) are shown. Mice were assigned to either stressor or control groups. Mice in social isolation were maintained alone in cages for a week (blue). Mice in the Int-HFD group accessed a HFD for two hours during the day every other day for two weeks (orange). For the restraint group, 7-week-old mice were subjected to two hours of complete immobilization using restrainers for five consecutive days (green). Yellow boxes indicate the day when the cannulas were placed into the NAcc of mice for microdialysis experiments. ***B***, Experimental scheme to assess mouse feeding behaviors. Four bait containers were affixed at 10 cm intervals along the outer edge of half a circle plate. The respective diet containers required mice to physically insert their heads to consume the bait. Mouse movements was captured and analyzed using imaging software. Mice normally feed equivalently from multiple bait sources. In contrast, stress-affected mice exhibited “fixated feeding” with a clear preference for a specific bait source (spatial feeding pattern). We also analyzed temporal feeding behaviors with aberrant feeding states characterized by “protracted feeding”, an uninterrupted prolonged period of feeding (temporal feeding pattern). ***C***, The spatial feeding patterns of mice reared in social isolation or in group housing (control). Representative heatmap images indicate the duration of time spent on a bait source (top images). The duration on each bait source was quantified by the motion capture system with the sources ranked in decreasing order in terms of duration on each bait position and a preference ratio was determined (bottom graph). Similarly, the amount consumed from each source was quantified with the sources ranked in decreasing order in terms of bait consumption (Fig. 1-1). Statistical analysis was performed on the preference ratio (n = 15 for isolation, n = 16 for control, Welch’s t test). ***D***, The spatial feeding patterns of mice with the Int-HFD diet or a normal diet (control). Representative heatmap images (top images) reflect the duration of time spent on a bait source as before. The duration on each bait source was quantified by the motion capture system (bottom graph) and statistical analysis were performed as in *C* (n = 16 for each, Welch’s t test). ***E***, The spatial feeding patterns of physically restrained and non-restrained mice (control). Representative heatmap images (top images), quantification of the duration of time spent on each source (bottom graph), and statistical analysis were performed as in *C* (n = 10 for restrained, n = 11for control, Welch’s t test). \**P* < 0.05. Data are mean ± SEM.

Dopamine transporter (DAT)-Cre mice (B6.SJL-*Slc6a3*^*tm1*.*1(cre)Bkmn*^/J) were obtained from Jackson Laboratories. The tyrosine hydroxylase transporter (TH)-Cre mice (B6.FVB(Cg)-Tg(Th-cre)FI172Gsat/Mmucd) were obtained from MMRRC.

All animal experiments were performed in accordance with the National Institute of Health Guide for the Care and Use of Laboratory Animals and were approved by the Nagoya University Animal Experiment Committee. Water was provided ad libitum.

### Real-time monitoring system

To eliminate any potential artificial effects on the reward system, we developed a real-time monitoring system in tandem with regular diet instead of palatable foods and operant conditions to assess both the spatial and temporal aspects of feeding behaviors. This approach allows for more accurate detection of not only the relative physiological states of animals fed regular diet but also subtle alterations in feeding behaviors.

The real-time monitoring system consists of a box containing standard chow diet (CE2, CLEA) affixed to the surface of a 60-cm diameter circle originally designed for open field tests. Four bait containers (33 × 37 × 22 mm, SHINFACTORY, Fukuoka, Japan) were affixed at 10-cm intervals along the outer edge of one half of the circle plate in close proximity to the wall (height, 40 cm) (Fig. 1*B*, left). The respective diet containers were designed such that the mice needed to physically insert their heads into the container to consume the bait. This movement was captured and analyzed by the motion capture system using EthoVision XT 12 (Noldus Information Technology, Wageningen, the Netherlands) imaging software. The software was used to monitor approaches to bait containers, the duration of each feeding movement in the arena. Prior to the start of the experiment, mice were deprived of a food source for 4 h, after which their feeding behavior was observed for 4 h using the real-time monitoring system. At the conclusion of the experiment, the residual bait in each container was weighed.

### Definitions of fixated and protracted feeding

Using the real-time monitoring system, we defined two distinct aberrant feeding behavior pattern phenotypes -“fixated feeding” and “protracted feeding” (Fig. *1B*, right). Fixated feeding is characterized by an abnormal deviation of food selection. For instance, mice with spatial fixation approached a specific bait container more frequently than the other containers, whereas control mice had no bait preference. Protracted feeding is characterized by a prolonged uninterrupted feeding interval, such that mice with this phenotype spent significantly longer periods of time on a single eating event than control mice.

### Kinetic measurement of dopamine levels by in vivo microdialysis

The amount of dopamine in the NAcc shell was measured by microdialysis. The guide cannula was stereotaxically placed into the right side of the mouse NAcc shell (L: +0.42, A: -1.34, H: -3.95) a week prior to the measurement. Experimental mice were deprived of food for 15 h prior to the start of the assay. Dopamine level was monitored by HPLC-ECD (HTEC-510, Eicom, Kyoto, Japan) both before and after feeding. Dialysis probe (FX-6-01, Eicom) placements were verified histologically at the ends of each experiment, and experimental data were excluded if the membrane portions of the dialysis probes lay outside the NAcc shell region.

### Selective dopamine administration into the NAcc shell

Mice under the three experimental conditions selectively had dopamine administered into the NAcc shell. The guide cannula was stereotaxically placed into the right side of the mouse NAcc shell (L: +0.42, A: +1.34, H: -4.25) 1 week prior to assessing feeding behavior (Fig. 2-1*A*). The dopamine solution was prepared as 3-hydroxytyramine hydrochloride: C_8_H_11_NO_2_-HCl (Tokyo Chemical Industry, Tokyo, Japan) diluted in solution buffer [0.5% ascorbic acid (ASA; Iwaki Seiyaku, Tokyo, Japan)/Ringer’s solution]. Dopamine administration was performed 5 min prior to feeding behavior analyses by injecting 0.5 μl dopamine solution (200 μg/μl) into the NAcc shell at a flow rate of 0.25 μl /min. The injection cannula was maintained in place post-injection for 3 min. For control mice, the same volume of 0.5% ASA/Ringer’s solution was injected into the NAcc shell.

**Figure 2.**
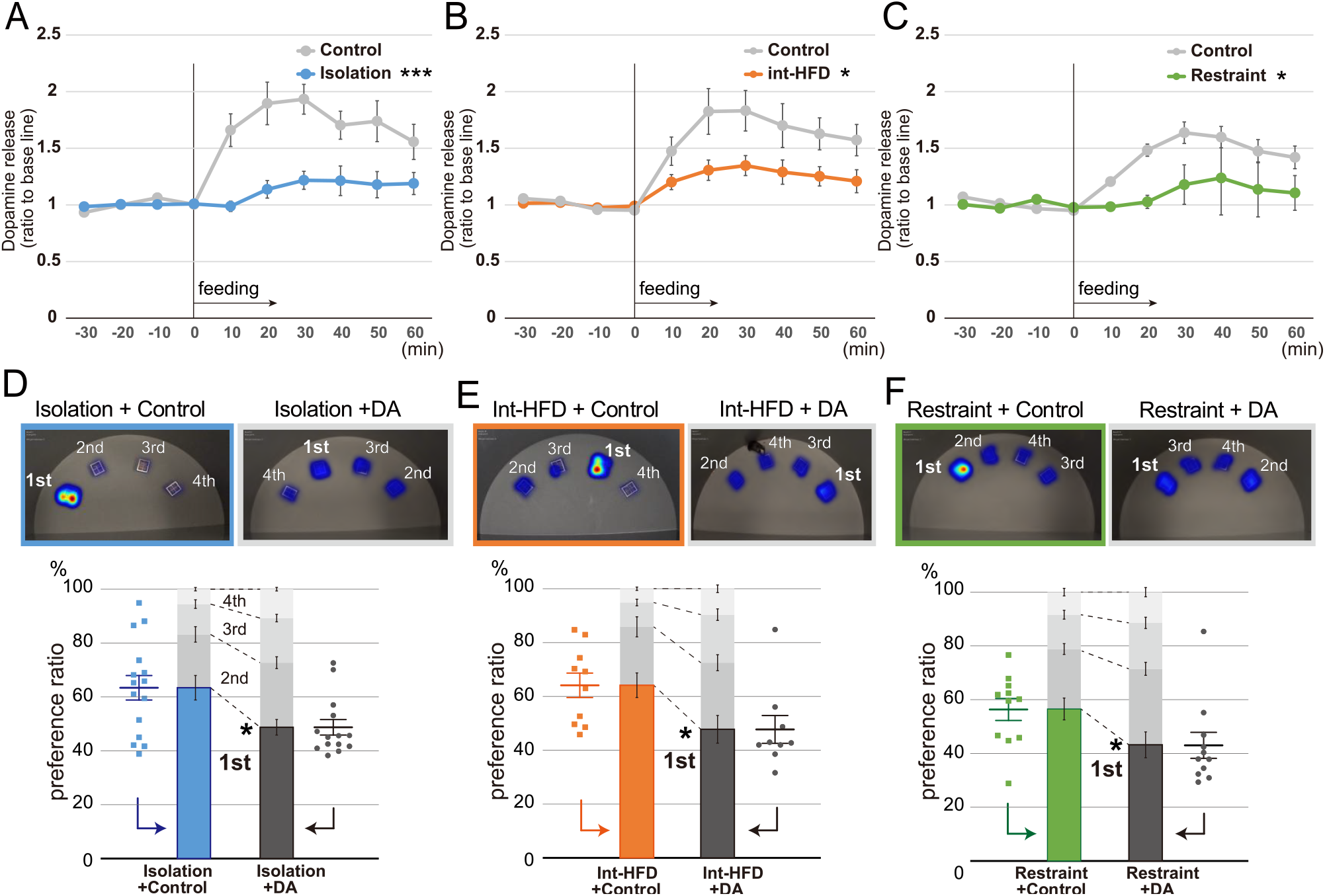
Dopamine levels in the NAcc shell is impaired by psychosocial stresses, whereas dopamine supplementation normalizes the stress-induced aberrant spatial feeding behavior patterns characterized by fixated feeding. ***A***, *In vivo* microdialysis analysis of mice from the isolation group. Feeding increased the amount of dopamine (DA) in the NAcc shell. The extracellular DA levels were determined by *in vivo* microdialysis and HPLC-ECD. Basal fractions were collected prior to the initiation of feeding at time point 0 (n = 8 each, ***significantly different from control group by ANOVA with repeated measures). ***B***, *In vivo* microdialysis analysis of mice from the Int-HFD group (n = 8 for Int-HFD, n=9 for control, *significantly different from control group by ANOVA with repeated measures). ***C***, *In vivo* microdialysis analysis of mice from the restrained group (n = 8 for restrained, n = 9 for control, *significantly different from control group by ANOVA with repeated measures). ***D***, The spatial feeding patterns of mice in the isolation + control groups and isolation + DA groups. Representative heatmap images (top images) are as in Fig. 1*C*. The duration of time spent per bait position was quantified by the motion capture system with the sources ranked as in Fig. 1c. Similarly, the amount consumed for each source were ranked in decreasing order (Fig. 2-1). Statistical analysis was performed as in Fig. 1*C* (n = 15 for isolation + control, n = 14 for isolation + DA, Mann-Whitney U test). ***E***, The spatial feeding patterns of mice in the Int-HFD + control and the Int-HFD + DA groups. Representative heatmap images (top images), graphs (bottom), and statistical analysis are as above (n = 10 for Int-HFD + control, n = 9 for Int-HFD + DA, Mann-Whitney U test). ***F***, The spatial feeding patterns of mice in the restraint + control and restraint + DA groups. Representative heatmap images (top images), graphs (bottom), and statistical analysis are as above (n = 11 for each, Mann-Whitney U test). \**P* < 0.05, \*\*\**P* < 0.001. Data are mean ± SEM.

### DREADD system

To investigate the functional properties of the dopaminergic neurons in the ventral tegmental area (VTA), we used a DREADD system employing AAVs expressing hM4Di with subsequent Clozapine N-oxide (CNO, R&D Systems, Minneapolis, MN) administration (Fig 3*A*). We injected 0.5 μl of AAV-mCherry-FLEX-hM4Di virus (2.0 × 10^13^ copies/ml) into the bilateral VTA regions (L: +/-0.4, A: -3.28, H: -4.50) of 6-week-old DAT-Cre mice and TH-Cre mice at a flow rate of 0.25 μl /min. The injection cannula was maintained in place post-injection for 3 min. CNO [1.0 μg/g of BW, 0.1 μg/μl in 10% DMSO (Sigma-Aldrich, St. Louis, MO) -saline mixture] was intraperitoneally administered 30 min prior to real-time monitoring of feeding behavior or *in vivo* microdialysis. For controls, the same volume of 10% DMSO -saline mixture was administered intraperitoneally.

**Figure 3.**
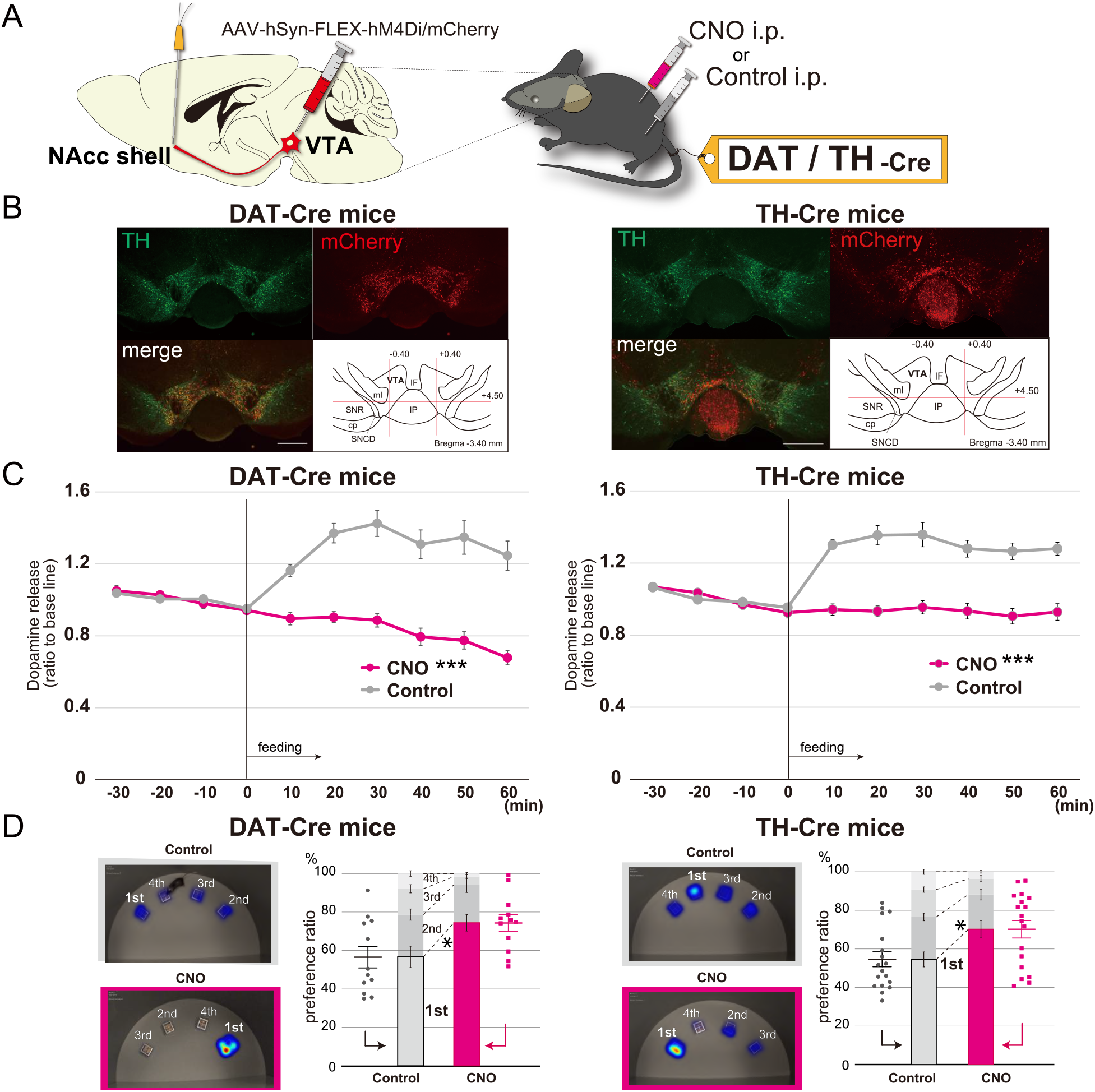
Selective inhibition of the dopaminergic neuronal circuit from the VTA to the NAcc replicates fixated feeding behavior patterns. ***A***, DREADD-based experimental scheme for inhibiting the dopaminergic neurons at the VTA. AAV-hSyn-FLEX-hM4Di/mCherry was injected into the VTA of 6-week-old DAT-Cre or TH-Cre mice (see also Fig. 3-1*A*). ***B***, Immunofluorescent images of dopaminergic neurons in the VTA with mCherry expression. Sections including the VTA were stained with anti-TH antibodies. Abbreviations used in the figure: cp = cerebral peduncle; IF = interfascicular nucleus, IP = interpeduncular nucleus, ml = medial lemniscus; SNCD = substantia nigra, compact part, dorsal tier; SNR = substantia nigra, reticular part. Scale bar, 100 μm. ***C***, *In vivo* microdialysis analysis of DAT-Cre (left) or TH-Cre (right) mice in which AAV-hSyn-FLEX-hM4Di/mCherry was injected into the VTA. Feeding after overnight starvation induced the release of DA in the NAcc shell. Mice were intraperitoneally administered CNO or control 30 min prior to feeding. The extracellular DA levels of mice were determined as in Fig. 2 (n = 8 for DAT-Cre mice in the left graph, n = 11 for TH-Cre mice in the right graph, ***significantly different from control groups by ANOVA with repeated measures). ***D***, The spatial feeding patterns of DAT-Cre mice (left) or TH-Cre mice (right) treated with CNO or control following AAV-hSyn-FLEX-hM4Di/mCherry injection. Representative heatmap images (left images) and preference ratios were determined as in Fig. 1*C* (see also Fig. 3-1*B*). Statistical analysis was performed on the most frequently consumed bait ratio (n = 12 for DAT-Cre mice, n = 18 for TH-Cre mice, Wilcoxon signed-rank test). \**P* < 0.05, \*\*\**P* < 0.001. Data are mean ± SEM.

### Body composition and tissue examination

Body weight, rectal temperature, and blood glucose were monitored in 8-week-old mice. Dissection and collection of tissue samples including brain, spleen, pancreas, white adipose tissue (WAT), and brown adipose tissue (BAT) were likewise performed using 8-week-old mice synchronized to the same time schedule as mice in the microdialysis or feeding behavior monitoring experiments. Mice were allowed ad libitum access to water and standard bait prior to dissection. Blood glucose levels were measured before anesthetization. After cooling on ice, the brain was sectioned with a 1-mm thick brain slicer (MBS-A1C, BrainScience idea, Osaka, Japan) and the hypothalamus was collected under a stereomicroscope and rapidly cooled to -80°C along with the WAT and BAT. Rectal temperature measurements and plasma catecholamine concentration determinations were performed immediately after each final stress under anesthesia. Rectal temperatures were measured by inserting a probe (TX10-01, 900-21B, Kenis, Osaka, Japan) coated with mineral oil (Sigma-Aldrich) into the anus and fixing it at a distance of 2-cm, and then measuring the temperature after stabilization. The plasma catecholamine concentration was measured by intracardiac blood sampling with samples immediately placed in EDTA-2K anticoagulant spits (BD Japan, Tokyo, Japan) and centrifuged at 1500 x g for 10 min. The supernatant was collected and passed through a clean EG column (CA-50 PBA, Eicom, Kyoto, Japan) prior to sequential HPLC-ECD (HTEC-510, Eicom) loading to determine the catecholamine concentration. 3,4-Dihydroxybenzylamine (858781, Sigma-Aldrich) was used as an internal standard.

### Immunohistochemistry

Mouse brains were fixed in 4% paraformaldehyde and cut into 40-μm-thick sections using a vibratome (Leica Biosystems, Nussloch, Germany). The sections were blocked using blocking solution (X0909, Agilent Technologies, Santa Clara, CA) for 1 h followed by overnight incubation with anti-TH antibody (AB152, Sigma-Aldrich) diluted in antibody diluent (S0809, Agilent Technologies) at 4°C. For immunofluorescence, Alexa-488-conjugated secondary antibody (Thermo Fisher Scientific, Waltham, MA) and DAPI were used.

### Real-time qPCR for gene expression analysis

An RNeasy Mini Kit (Qiagen, Hilden, Germany) was used to isolate total RNA from tissues. cDNA synthesis was subsequently performed using 1 μg total RNA and oligo-dT primers (C110A, Promega, Madison, WI). Transcript levels were assessed via real-time quantitative polymerase chain reaction (qPCR) amplifications using an iQ SYBR Green Supermix (BioRad, Hercules, CA) with primers listed in Table 4-1 on a CFX96 system (BioRad). Thermocycler conditions were 95°C for 3 min followed by 40 cycles of 95°C for 10 sec and 55°C for 30 sec. The relative quantity of each transcript was determined via a standard curve using the cycle thresholds of cDNA serial dilutions normalized to *Gapdh* levels. PCR was performed in triplicate for each sample, and all experiments were replicated twice.

### Statistical analysis

Statistical analyses were performed using JMP 15 (SAS Institute, Drive Cary, NC). The Anderson-Darling test was used to test normality of samples in two groups. The Welch’s t-test or paired t-test was used for samples with normal distribution, whereas Mann-Whitney U test or Wilcoxon signed-rank test was used for those without normal distribution as described in the Figure legends with statistical significance set to P < 0.05. Significant differences in groups were determined by ANOVA with repeated measures as indicated in the Figure legends of *in vivo* microdialysis. The data are expressed as mean ± SEM. All statistical tests were two-sided.

## Results

### Psychosocial stresses cause aberrant feeding behavior patterns characterized by fixated feeding

To investigate how feeding behavior patterns are regulated, we used a mouse model incorporating three different psychosocial stressors: social isolation, a major inducer of anxiety; intermittent high-fat-diet (HFD), which provokes dissatisfaction; and physical restraint, which mimics the physical stress caused by confinement (Fig. 1*A*). Both stress conditions and neuropsychological disorders can induce aberrant feeding behavior patterns such as fixations on one kind of food or overeating regardless of energy demands. To detect these aberrations, we examined impaired feeding states in terms of spatial and temporal feeding behavior patterns defined as “fixated feeding”, a preference for a particular bait, and “protracted feeding”, a prolonged uninterrupted feeding interval (Fig. 1*B*). Based on this experimental approach, we investigated aberrant feeding behavior patterns provoked in mice by psychosocial stressors.

By calculating the duration of time spent per bait position and the amount of bait consumed, we found that socially isolated mice fixated on a specific bait location. In contrast, feeding by control mice was more evenly distributed across the different bait positions (Fig. 1*C* and Fig. 1-1*A*). Similar fixation on a specific bait location was also observed in mice from the intermittent HFD and physically restrained groups (Fig. 1*D* and *E*, and Fig.1-1*B* and *C*), indicating a role for stress in provoking the fixated feeding behavior.

### Dopamine release in the NAcc shell is impaired by psychosocial stresses, whereas dopamine supplementation normalizes the stress-induced aberrant spatial feeding behavior patterns characterized by fixated feeding

To investigate the effects of stresses on the reward system, *in vivo* microdialysis analyses were performed to assess the amount of dopamine in the nucleus accumbens (NAcc) shell after feeding on a standard chow diet under all three stress conditions. In the social isolation study, mice from the control group exhibited an increase in dopamine levels over the initial amount at 10-30 min post-feeding. This dopamine peak was not observed in the socially isolated mice and the degree of dopamine increase was significantly lower in this group compared to the control group after feeding (Fig. 2*A*). Similar to the socially isolated mice, the increase in dopamine levels observed in the intermittent HFD and restraint mice was significantly reduced after feeding as compared to controls (Fig. 2*B* and *C*). These results indicate that psychosocial stresses inhibit the responsiveness of dopamine in the NAcc shell after feeding.

To confirm the relationship between aberrant dopamine levels in the NAcc shell and the impaired feeding behavior patterns, we selectively administered dopamine into the NAcc shell of mice under all three experimental conditions. Feeding behavior monitoring was performed after dopamine administration (Fig. 2-1*A*). Dopamine administration reversed fixation on the preferred bait position that had been observed in all three conditions (Fig. 2*D-F*, and Fig. 2-1*B-D*).

### Selective inhibition of the neuronal circuit from the VTA to the NAcc replicates fixated feeding patterns

The dopaminergic neurotransmission circuit from the ventral tegmental area (VTA) to the NAcc shell is largely associated with the reward/motivation system (Arias-Carrion et al., 2010). To investigate the relationship between this dopaminergic circuit and the aberrant feeding behavior patterns, we used the Designer Receptors Exclusively Activated by Designer Drugs (DREADD) system to selectively modulate the dopaminergic neurons at the VTA by incorporating AAVs expressing hM4Di with the FLEX conditional gene expression system in dopamine transporter-Cre transgenic (DAT-Cre) mice and tyrosine hydroxylase-Cre (TH-Cre) transgenic mice (Fig. 3*A* and Fig. 3-1*A*). After selectively inhibiting dopaminergic neuron activity at the VTA in both mice lines (Fig. 3*B*), we assessed the amount of dopamine levels at the NAcc shell. In response to feeding, dopamine levels increased when administered control solution, but was negligible when they were given clozapine N-oxide (CNO) (Fig. 3*C*). In addition, the inhibition of dopaminergic neuron activity from the VTA to NAcc shell resulted in a state of fixated feeding in both the DAT-Cre and TH-Cre mice, which indicates that the mesolimbic dopamine-mediated reward system is directly associated with aberrant feeding behavior patterns that are characterized by fixated feeding (Fig. 3*D* and Fig. 3-1*B*) To exclude nonspecific effects of CNO on spatial feeding behavior patterns, the spatial feeding patterns of mice treated with CNO or DMSO were also examined. Neither the duration of time spent per bait position or the amount consumed from each source was altered by the administration of CNO (Fig. 3-1*C*).

### The three psychosocial stressors had inconsistent effects on metabolic states

We investigated three different mouse models exhibiting aberrant feeding behaviors - social isolation, intermittent HFD treatment, and physical restraint on metabolic factors. Body weight and temperature, blood sugar, plasma norepinephrine concentration, the amounts of brown (BAT) and white adipose tissue (WAT), and spleen weight were investigated and we found that three psychosocial stressors had variable effects on the general metabolic factors (Fig. 4). Gene expression states in the hypothalamus, brown adipose tissue (BAT), and white adipose tissue (WAT) were also inconsistent among the stressors (Fig. 4-1). Since all of the psychosocial stressors promoted feeding patterns characterized by a state of fixated feeding, these findings indicate that the metabolic alterations induced by the varied psychosocial stress conditions are not direct causes of the aberrant feeding behavior patterns.

**Figure 4.**
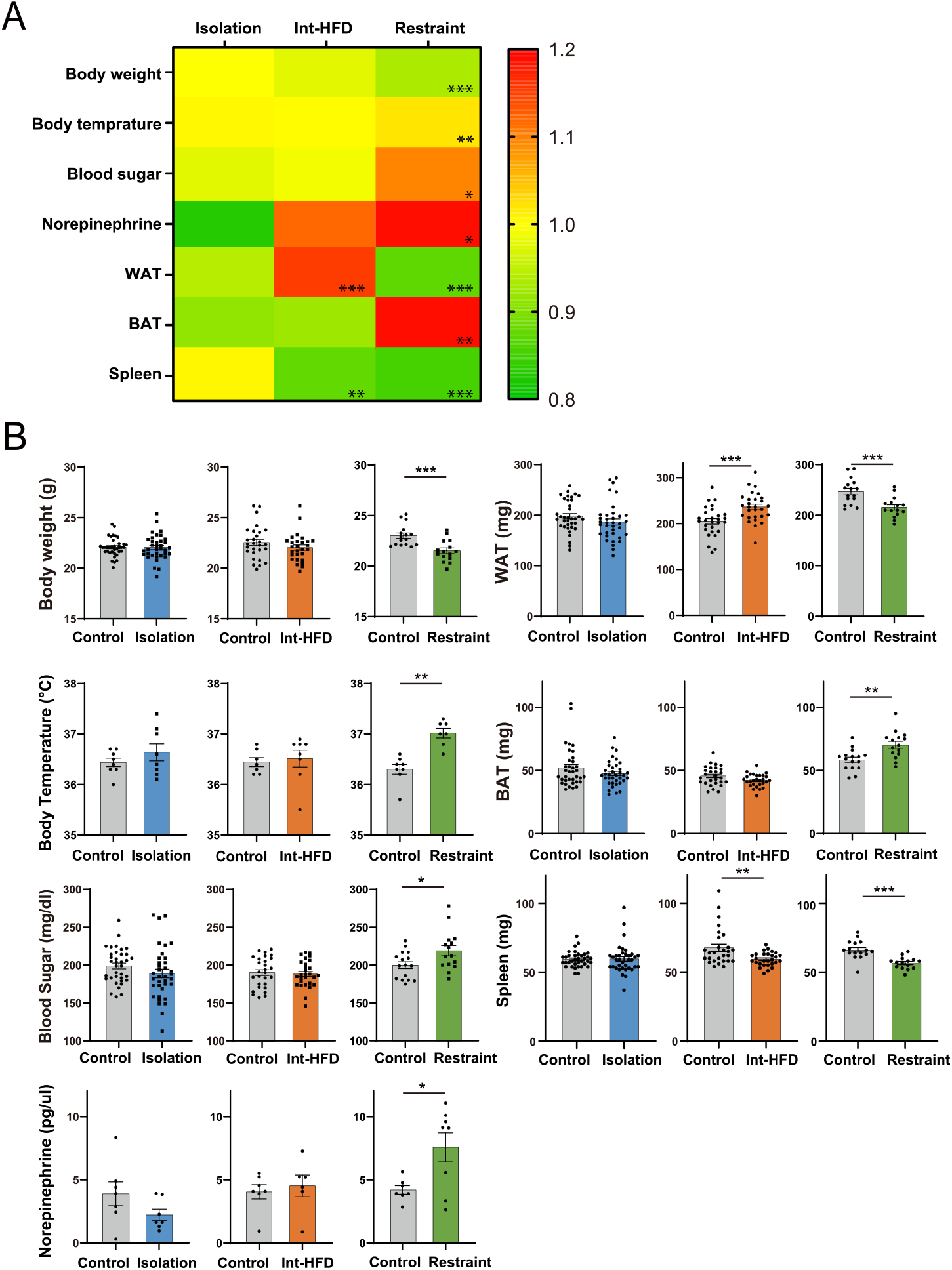
General characterization of metabolic factors in the three stressor models. ***A***, Effects of the three stressor conditions (social isolation, Int-HFD, and physical restraint) on metabolic factors. Body weight and temperature (rectal temperature), blood sugar, plasma norepinephrine concentration, the amounts of brown (BAT) and white adipose tissue (WAT), and spleen weight were determined. Results of the measurements across the experimental conditions are depicted via a heatmap (ratio to controls). Note that the norepinephrine data (isolation and restrained) are off scale. ***B***, Quantitative data are shown as comparisons between each respective stress condition (isolation, blue; Int-HFD, orange; restrained, green) and controls (Isolation group body temperature - n = 8 for both; Int-HFD group body temperature - n = 8 for Int-HFD, n = 7 for Control; Restrained body temperature - n = 7 for Restraint, n = 8 for Control; Mann-Whitney U test. Isolation group plasma norepinephrine - n = 7 for both; Int-HFD group plasma norepinephrine - n = 6 for Int-HFD, n = 7 for Control; Restrained plasma norepinephrine - n = 8 for Restraint, n = 7 for Control; Welch’s t test. Isolation group others - n = 36 for both; Int-HFD group others - n = 28 for both; Restrained group others - n = 15 for Restraint, n = 16 for Control; Mann-Whitney U test). *P < 0.05, **P < 0.01, ***P < 0.001. Data are mean ± SEM.

### Stressors other than restraint exhibit aberrant temporal feeding patterns via the mesolimbic dopamine system

In addition to the spatial feeding patterns, the temporal feeding patterns of mice were investigated in terms of feeding duration and food contact. The average duration of each feeding behavior was extended in mice from the social isolation and intermittent HFD groups relative to control mice, but was not affected in the physical restraint group (Fig. 5*A*). Dopamine administration significantly reduced the average duration of each feeding bout in mice from the social isolation and intermittent HFD groups (Fig. 5*B*). Moreover, inhibiting dopaminergic neuron activity from the VTA to the NAcc shell triggered a significant increase in the average duration of feeding in both the DAT-Cre and TH-Cre mice (Fig. 5*C*). To exclude nonspecific effects of CNO on spatial feeding behavior patterns, the temporal feeding patterns of mice treated with CNO or DMSO were also examined. The average duration of each feeding bout was not altered by the administration of CNO (Fig. 5-1).

**Figure 5.**
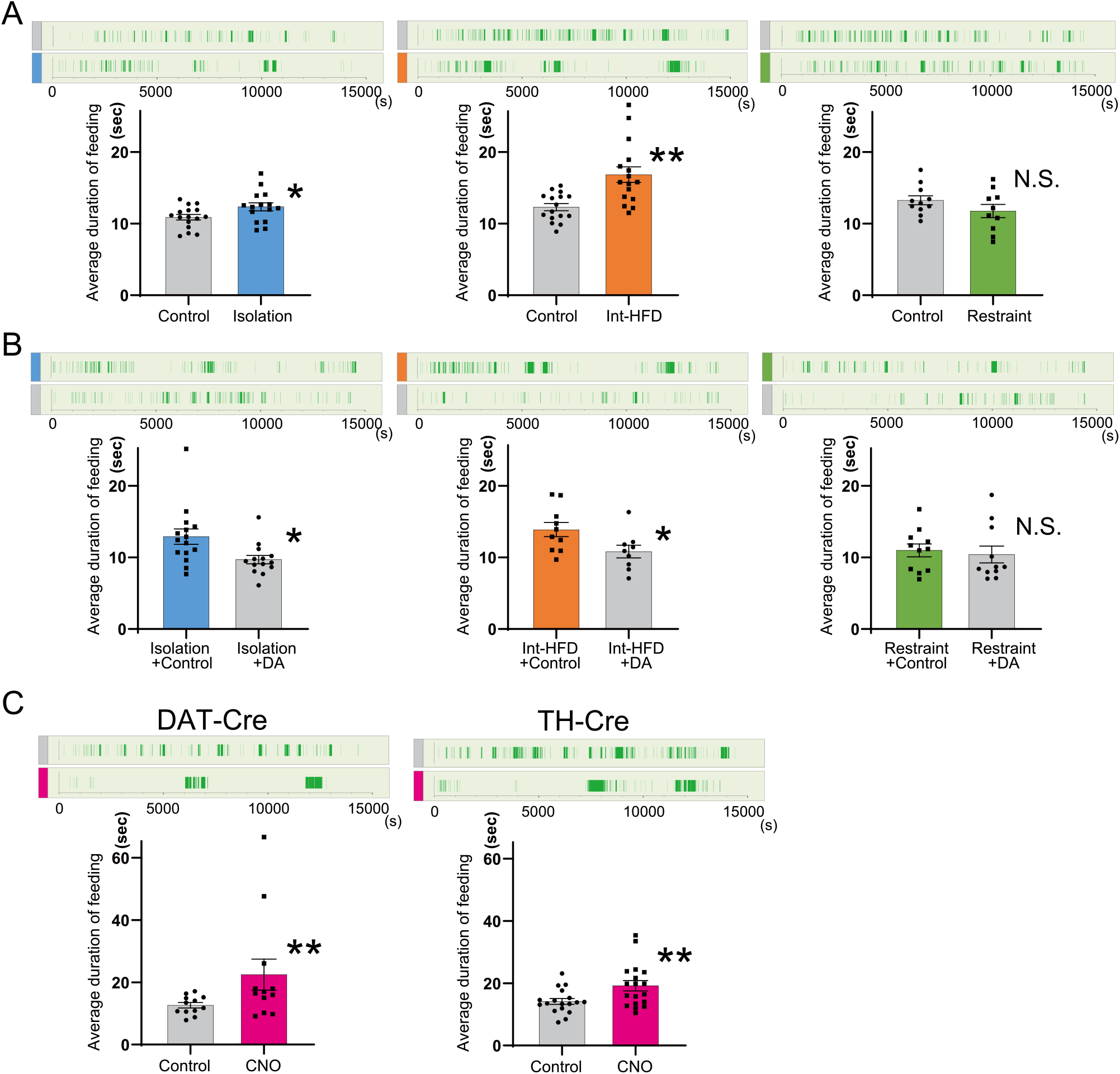
Stressors other than physical restraint exhibit aberrant temporal feeding behavior patterns that are characterized by protracted feeding via the mesolimbic dopamine system. ***A***, The temporal feeding patterns of mice reared in isolation or group housing, mice fed Int-HFD or a normal diet, and physically restrained or non-restrained mice were investigated. Representative graphs of feeding duration over the course of the session are shown (top graphs, dark green indicates feeding duration). The average durations of feeding were compared between the stress and control groups (left panel - isolation group, n = 15 for isolation, n = 16 for control; middle panel - Int-HFD group, n = 16 for each; right panel - restrained group, n =10 for restrained, n = 11 for control, Welch’s t test). ***B***, The temporal feeding patterns of mice in the isolation + control and isolation + DA groups, the Int-HFD + control and Int-HFD + DA groups, and the restraint + control and restraint + DA groups were investigated as well (left panel - isolation rescue, n = 15 for isolation + control, n = 14 for isolation + DA; middle panel - Int-HFD rescue, n = 10 for int-HFD + control, n = 9 for Int-HFD + DA; right panel - restrained rescue, n = 11 for each, Mann-Whitney U test). ***C***, The temporal feeding patterns of DAT-Cre mice (left panel) or TH-Cre mice (right panel) treated with CNO or control following AAV-hSyn-FLEX-hM4Di/mCherry injection were examined (n = 12 for DAT-Cre mice, n = 18 for TH-Cre mice, Wilcoxon signed-rank test). \**P* < 0.05, \*\**P* < 0.01, N.S. denotes not significant. Data are mean ± SEM.

## Discussion

We found that three different external stressors, all of which altered the feeding patterns of mice, deviations in food selection (fixated feeding) regardless of alterations in metabolic conditions. These findings indicate that feeding behaviors can be quantified not only by total intake but also by variations in spatial patterns that enable the detection of psychosocial stressor-mediated changes in the reward system as well as the prodromal state of neuropsychiatric diseases. Moreover, the mesolimbic dopamine-mediated reward system also contributes to aberrant temporal feeding behavior patterns characterized by protracted feeding. The lack of a protracted feeding response in the restrained stress mice suggests the presence of other factors that obscure the effects of the reward system on regulation of temporal feeding patterns (Fig. 5).

Physical restraint stress excites sympathetic neurons, leading to increased plasma norepinephrine, increased blood sugar, increased body temperature, decreased spleen weight, and increased BAT, all of which were observed in our restraint mice but not in the other two stressor models (Fig. 4*A*). Thus, it was assumed that excitation of sympathetic neurons may impact the temporal feeding pattern.

Intermittent HFD treatment is frequently used as a model for binge eating, which can be caused by various psychosocial stresses (Novelle and Dieguez, 2018). Because we sought to detect early traces of feeding behavior alterations, our external stimuli application period, including intermittent HFD treatment, was limited to a relatively short duration (1-2 weeks). Accordingly, we found that mice in the intermittent HFD group exhibited fixated feeding pattern, but maintained their body weight despite an increase in total food intake per session (data not shown) (Rospond et al., 2015). Conversely, while physical restraint stress also showed a similar fixated feeding pattern, it provoked a significant but marginal decrease in body weight (Fig. 4). As stressors can induce both weight gain and loss in humans, the three psychosocial stressors examined in our mouse model had inconsistent effects on both the amount of bait consumed and body weight. In contrast, they consistently induced fixated feeding, indicating that alterations in feeding behavior patterns precede changes in body weight under psychosocial stress conditions.

We determined that dopamine neurons from the VTA to the NAcc shell participate in generating spatiotemporal feeding behavior patterns. Food intake rapidly evokes dopaminergic neuron activity in the VTA, which leads to dopamine release in the NAcc (Gunaydin et al., 2014). A previous study revealed that mice under stress exhibited similar impaired dopamine responses following ethanol intake (Ostroumov et al., 2016). While activation of dopaminergic neurons in the VTA has been reported to suppress reward-driven food consumption (Adamantidis et al., 2011; Mikhailova et al., 2016), our results indicate that feeding behavior patterns are positively regulated by dopaminergic system from the VTA to the NAcc shell regardless of the amount of food intake. Considering that dopamine neurons in the VTA are the center of the reward system and that stresses impact dopaminergic neuronal activity in the mesolimbic dopamine system (Bloomfield et al., 2019), changes in spatiotemporal feeding behavior patterns may be indicative of impairments to the reward system that are induced by external stress.

These stress-induced alterations may also reflect innate escape responses in animals since integrated eating behaviors can be beneficial under potential environmental dangers such as predation. From this point of view, external stress induced changes in the reward system might be an adaptative response by the brain, similar to that seen in alterations in the hypothalamus or amygdala (Dietrich et al., 2015; Campos et al., 2018; Xu et al., 2019). Furthermore, the quality of feeding behaviors can be impacted by some forms of dementia, in particular FTLD as well as ASD (Ikeda et al., 2002; Ahmed et al., 2016; Bandini et al., 2017). Moreover, the mesolimbic reward pathway is affected by these neuropsychiatric diseases (Kloeters et al., 2013; Tye et al., 2013; Supekar et al., 2018). As such, altered feeding behavior patterns may be an early biomarker for a spectrum of neuropsychiatric diseases.

In summary, our study showed that psychosocial stress can rapidly alter spatiotemporal feeding behavior patterns via the limbic dopamine system. The detection and quantification of the spatiotemporal aspects of these feeding behavior patterns can be powerful tools for investigating the effects of various stressors such as those exacerbated by the recent COVID-19 pandemic.

## Abbreviations

AAV: Adeno-associated virus
ASA: ascorbic acid
ASD: autism spectrum disorders
BAT: brown adipose tissue
CNO: Clozapine N-oxide
COVID-19: coronavirus disease of 2019
cp: cerebral peduncle
DA: dopamine
DAPI: 4’,6-diamidino-2-phenylindole
DAT: Dopamine transporter
DMSO: Dimethyl sulfoxide
DREADD: Designer Receptors Exclusively Activated by Designer Drugs
EDTA: ethylenediaminetetraacetic acid
FLEX: flip-excision
FTLD: front temporal lobar degeneration
HFD: high-fat diet
hM4Di: human Gi-coupled M4 muscarinic receptor
HPLC-ECD: high performance liquid chromatography with electrochemical detector
hSyn: human synapsin
IF: interfascicular nucleus
Int-HFD: intermittent high fat diet
IP: interpeduncular nucleus
mCherry: Monomeric Cherry
ml: medial lemniscus
NAcc: Nucleus accumbens
qPCR: quantitative polymerase chain reaction
SEM: standard error of the mean
SNCD: substantia nigra, compact part, dorsal tier
SNR: substantia nigra, reticular part
TH: tyrosine hydroxylase transporter
VTA: ventral tegmental area
WAT: white adipose tissue

## Acknowledgements

Authors are grateful to the CARE (The Center for Animal Research and Education) at Nagoya University for technical support for the animal experiments and the Division of Experimental Animals at Nagoya University School of Medicine.

## Author contributions

Conceptualization, Y.F., S.I., and G.S.; Mice experiments, Y.F., K.K., K.E., N.I., M.I., D.T., and S.I.; Mice and AAV preparation, K.K., T.Y, and A.Y.; Data analysis, Y.F., K.K., M.K., H.W., and S.I.; Writing – Original Draft, S.I.; Writing – Review & Editing, Y.F., S.I., and G.S.; Funding Acquisition, S.I., and G.S.

