## Supplemental information for "Stress-impaired reward pathway promotes distinct feeding behavior patterns"

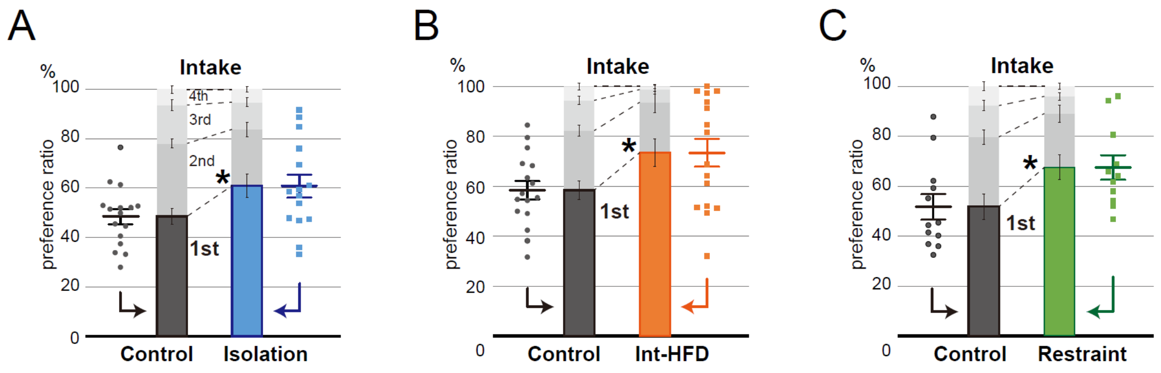


**Figure 1-1. Psychosocial stresses such as social isolation, intermittent high-fat diet, and physical restraint cause aberrant feeding behavior patterns characterized by fixated feeding.**
***A*****,** The spatial feeding patterns of mice reared in social isolation or group housing (n = 15 for isolation, n = 16 for control; Welch's t test). ***B*****,** The spatial feeding patterns of mice fed Int-HFD or a normal diet (n = 16 for both; Welch's t test). ***C*****,** The spatial feeding patterns of physically restrained and non-restrained mice (n = 11 for restraint, n = 12 for control; Welch's t test). In all cases, the amount consumed from each source was quantified with the sources ranked in decreasing order in terms of bait consumption. A preference ratio was determined and statistical analysis performed on the most frequently consumed bait ratio. $^*P < 0.05$. Data are mean ± SEM.

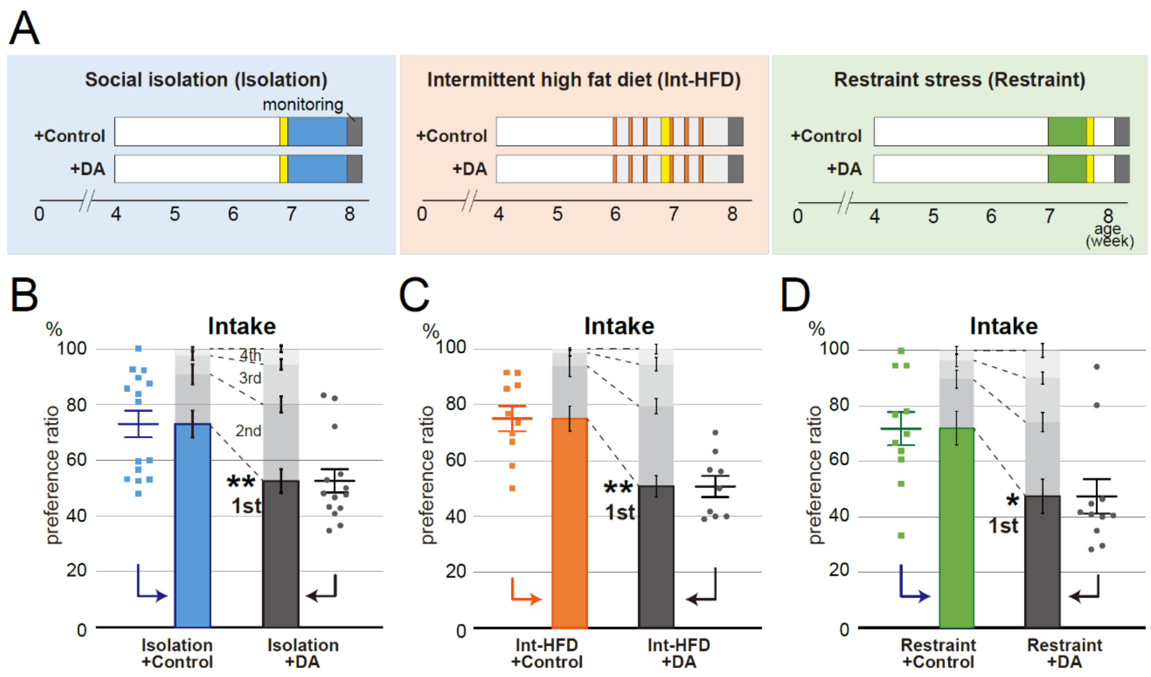


**Figure 2-1. Dopamine supplementation in the NAcc shell normalized the aberrant feeding behaviors characterized by fixative feeding.**

***A*,** Schematic of the study paradigm consisting of isolation + DA and isolation + control groups, Int-HFD + DA and Int-HFD + control groups, and restrained + DA and restrained + control groups. Mice were assigned into two groups, stressors with DA and stressors with 0.5% ASA/Ringer's solution (Control). After stress treatment, real-time feeding behavior with DA or control administration was monitored for each mouse after 4h starvation. Mice were treated with stressors as shown in Fig. 1A. Mice in the isolation group were housed alone in cages for a week (blue). Mice in the Int-HFD group were provided access to HFD for 2 h during the day every other day for 2 weeks (orange). For the restrained group, mice were immobilized with restrainers for 2 h over 5 consecutive days (green). Yellow boxes indicate the day when the cannulas were placed into the NAcc shell of mice. ***B*,** The spatial feeding patterns of mice in the isolation + DA and isolation + control groups (n = 15 for isolation + control, n = 14 for isolation + DA; Mann-Whitney U test). ***C*,** The spatial feeding patterns of mice in the Int-HFD + DA and Int-HFD + control groups (n = 10 for Int-HFD + control, n = 9 for Int-HFD + DA; Mann-Whitney U test). ***D*,** The spatial feeding patterns of mice in the restrained + DA and restrained + control groups (n = 11 for both; Mann-Whitney U test). For Fig. 2-1 *B-D*, the amount consumed from each source was quantified with the sources ranked in decreasing order in terms of bait consumption. A preference ratio was determined and statistical analysis was performed on the most frequently consumed bait ratio. $*P < 0.05$, $**P < 0.01$. Data are mean ± SEM.

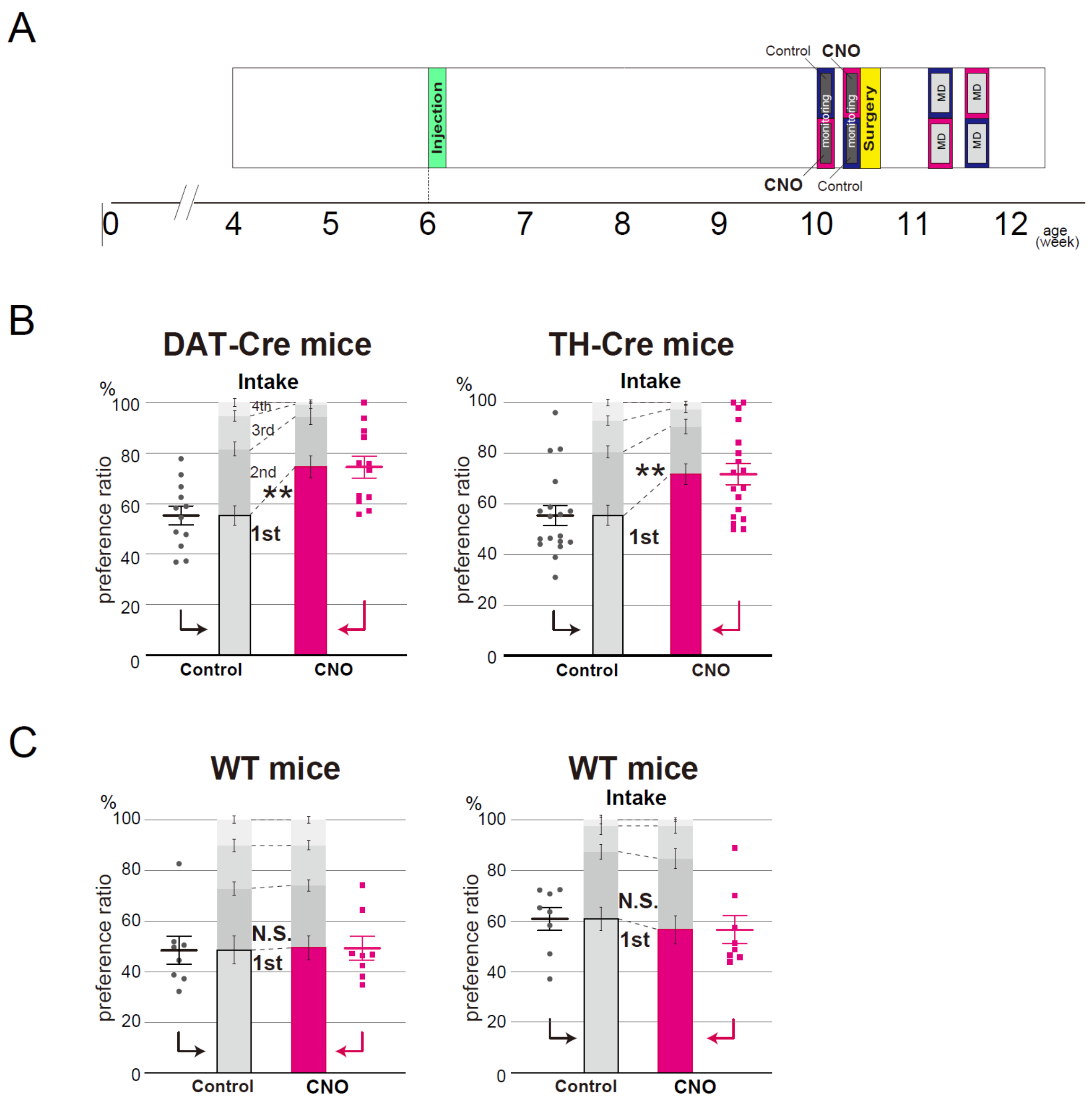


**Figure 3-1. Fixated spatial feeding behaviors are replicated by selectively inhibiting the dopaminergic neurons from the VTA to the NAcc shell**

***A*,** Schematic of the experimental design used to inhibit the dopaminergic neuronal circuit via the DREADD system. AAV-hSyn-FLEX-hM4Di/mCherry was injected into the VTA of 6-week-old DAT-Cre or TH-Cre mice (green boxes). The real-time feeding behavior of 10-week-old mice was monitored after 4h starvation. CNO or control were intraperitonially administered to inhibit the activity of hM4Di positive dopaminergic neurons in the VTA 30 min prior to behavioral monitoring. Dopamine release was measured by *in vivo* microdialysis when mice were 11-week-old. Yellow boxes indicate the day when the cannulas were placed into the NAcc shell of mice for the microdialysis experiments. ***B*,** The spatial feeding patterns of DAT-Cre mice (left) or TH-Cre mice (right) treated with CNO or control following AAV-hSyn-FLEX-hM4Di/mCherry injection. The amount consumed from each source was quantified with the sources ranked in decreasing order in terms of bait consumption. A preference ratio was determined and statistical analysis performed on the most frequently consumed bait ratio (n = 12 for each in DAT-Cre mice, n = 18 for each in TH-Cre mice; Wilcoxon signed-rank test). ***C*,** To exclude nonspecific effects of CNO on spatial feeding behavior patterns, mice treated with CNO or control were similarly examined. The duration of time spent per bait position was quantified by the motion capture system (left) and the amount consumed from each source was measured (right). No statistical differences were observed in the preference ratios between the CNO and control groups. Statistical analysis was performed on the most frequently consumed bait ratio ($n = 8$ for each; Wilcoxon signed-rank test). $^{**}P < 0.01$, N.S. denotes not significant. Data are mean ± SEM.

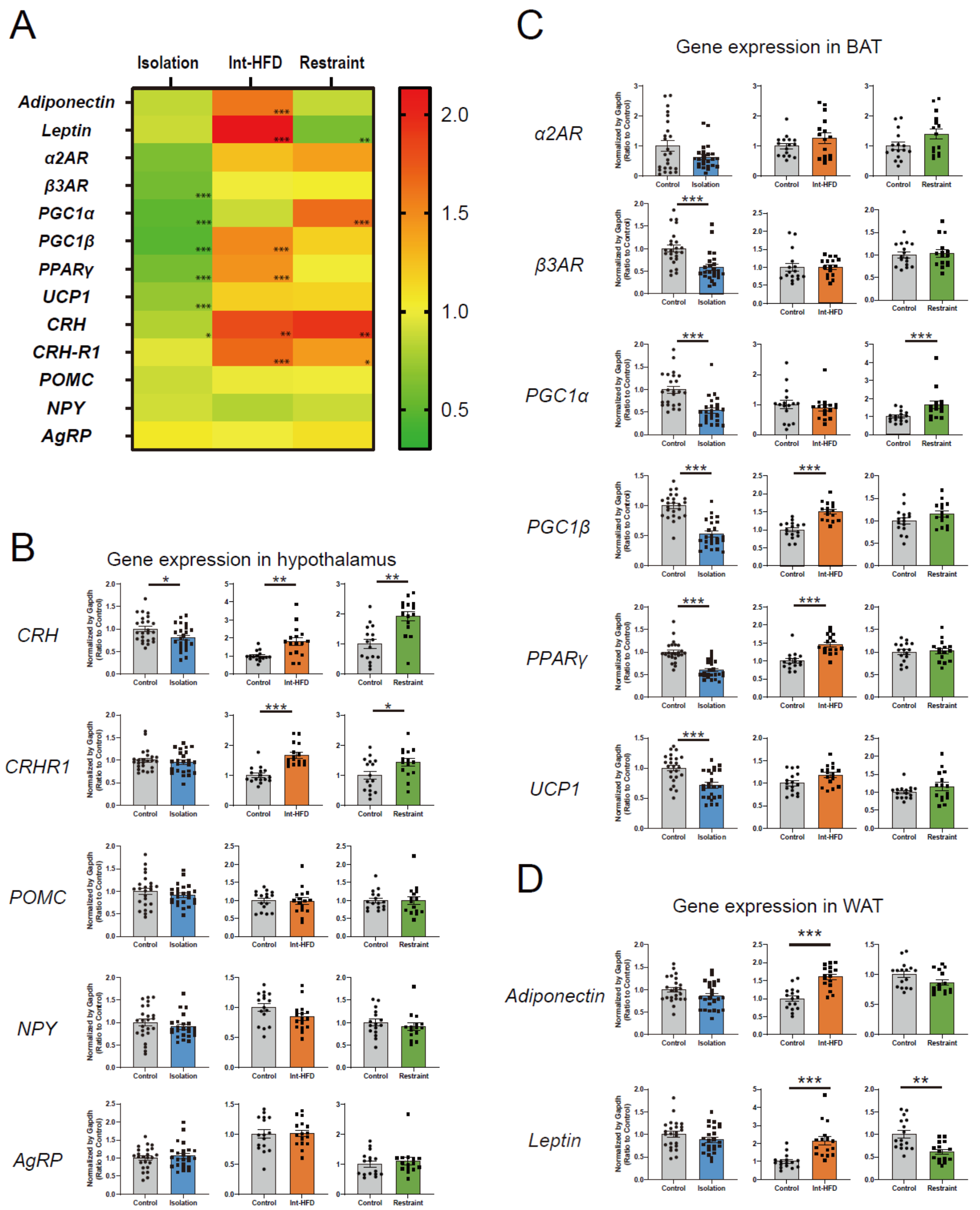


**Figure 4-1. Alterations in hypothalamus, BAT, and WAT-associated gene expression in the three stressor models.**

***A*,** Alterations of gene expression in the hypothalamus, BAT, and WAT under the three stress conditions (social isolation, Int-HFD, and physical restraint). The expression of *Adiponectin*, *Leptin*, *α2AR*, *β3AR*, *PGC1α*, *PGC1β*, *PPARγ*, *UCP1*, *CRH*, *CRH-R1*, *POMC*, *NPY*, and *AgRP* were measured by qPCR. The relative amount for each transcript was normalized to the amount of the house-keeping *Gapdh* levels. The heatmap summarizes the combined data. ***B*,** qPCR-based analyses of *CRH*, *CRH-R1*, *POMC*, *NPY*, and *AgRP* gene expression in the hypothalamus. Quantitative data are shown as a comparison between stressors (isolation, blue; Int-HFD, orange; restrained, green) and controls (isolation group - n = 24 for both; Int-HFD group - n = 16 for both; restrained group - n = 15 for restrained mice, n = 16 for control mice; Mann-Whitney U test). ***C*, *D*,** Expression levels of genes in BAT (C: *α2AR*, *β3AR*, *PGC1α*, *PGC1β*, *PPARγ*, and *UCP1*) and WAT (D: *Adiponectin* and *Leptin*). Quantitative data are shown as a comparison between stressors (isolation, blue; Int-HFD, orange; restrained, green) and controls (isolation group - n = 24 for both; Int-HFD group - n = 16 for both; restrained group - n = 15 for restrained, n = 16 for control; Mann-Whitney U test). $^{*}P < 0.05$, $^{**}P < 0.01$, $^{***}P < 0.001$. Data are mean ± SEM.

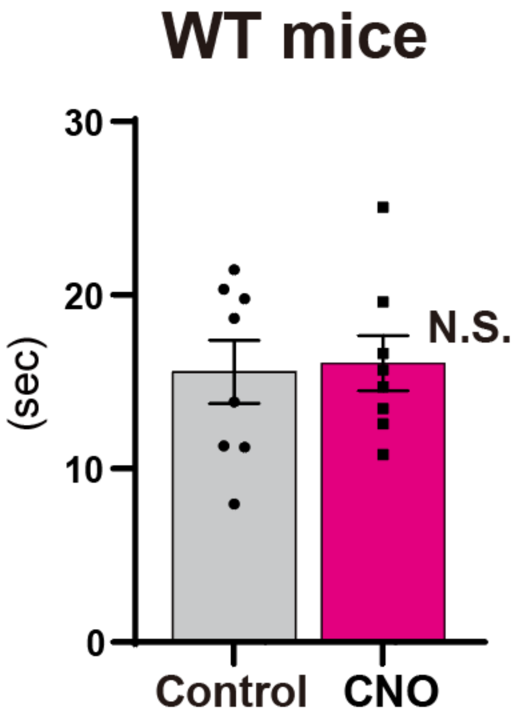


**Figure 5-1. Temporal feeding behaviors are not impacted by the administration of CNO in mice.** The temporal feeding patterns of mice treated with CNO or control were examined. The average durations of feeding were compared between the two groups (n = 8 for each; paired t-test). There was no statistical difference between the CNO and control groups. N.S. denotes not significant. Data are mean ± SEM.

**Table 4-1.**

**Primers used for qPCR in Figure 4-1.**

| | Forward | Reverse |
|---|---|---|
| **WAT** | | |
| *Adiponectin* | GCACTGGCAAGTTCTACTGCAA | GTAGGTGAAGAGAACGGCCTTGT |
| *Leptin* | GCCAGGCTGCCAGAATTG | CTGCCCCCCAGTTTGATG |
| **BAT** | | |
| *α2A-AR* | GCTGGTTATTATCGCGGTGT | CAGCGCCCTTCTTCTCTATG |
| *β3-AR* | CCTTCCGTCGTCTTCTGTGT | CAGCTTCCTTGCTGGATCTT |
| *PGC-1α* | CCCTGCCATTGTTAAGACC | TGCTGCTGTTCCTGTTTTC |
| *PGC-1β* | TCCTGTAAAAGCCCGGAGTAT | GCTCTGGTAGGGGCAGTGA |
| *PPAR γ* | GTGCCAGTTTCGATCCGTAGA | GGCCAGCATCGTGTAGATGA |
| *UCP1* | ACTGCCACACCTCCAGTCATT | CTTTGCCTCACTCAGGATTGG |
| **hypothalamus** | | |
| *CRH* | TCTGCGGGAAGTCTTGGAAA | TCCTGTTGCTGTGAGCTTGCT |
| *CRH-R1* | TGTTCGGTGAGGGCTGCTA | CGGTCGGTGGAGTACGTGAGT |
| *POMC* | CTGCTTCAGACCTCCATAGATGTG | CAGCGAGAGGTCGAGTTTGC |
| *NPY* | TCAGACCTCTTAATGAAGGAAAGCA | GAGAACAAGTTTCATTTCCCATCA |
| *AgRP* | CAGAAGCTTTGGCGGAGGT | AGGACTCGTGCAGCCTTACAC |
| *Gapdh* | TGTCGTGGAGTCTACTGGTG | GGCGGAGATGATGACCCTTT |

Abbreviations used in the Table 4-1: AgRP = agouti-related protein; α2AR = α2-adrenergic receptor; β3AR = β3-adrenergic receptor; CRH = corticotropin-releasing hormone; CRH-R1 = corticotropin releasing hormone receptor 1; NPY = neuropeptide Y; PGC1α = peroxisome proliferator-activated receptor gamma coactivator 1-alpha; PGC1β = peroxisome proliferator-activated receptor gamma coactivator 1-beta; POMC = proopiomelanocortin; PPARγ = peroxisome proliferator-activated receptor gamma; UCP1 = mitochondrial uncoupling protein 1
